# Capsidiol, a defensive sesquiterpene produced by wild tobacco in response to attack from the fungal pathogen *Alternaria alternata*, is regulated by an ERF2-like transcription factor

**DOI:** 10.1101/573675

**Authors:** Na Song, Lan Ma, Weiguang Wang, Huanhuan Sun, Lei Wang, Ian T. Baldwin, Jinsong Wu

## Abstract

Capsidiol is a sesquiterpenoid phytoalexin produced in *Nicotiana* and *Capsicum* species in response to pathogen attack. Whether capsidiol plays a defensive role and how its biosynthesis is regulated in the wild tobacco *Nicotiana attenuata* when the plant is attacked by *Alternaria alternata* (tobacco pathotype), a notorious necrotrophic fungus causing brown spot disease, is unknown. Transcriptome analysis indicated that a metabolic switch to sesquiterpene biosynthesis occurred in young leaves of *N. attenuata* after *A. alternata* inoculation: many genes leading to sesquiterpene production were strongly up-regulated, including the capsidiol biosynthetic genes, 5-*epi*-*aristolochene synthase* (*EAS*) and 5-*epi*-*aristolochene hydroxylase* (*EAH*). Consistently, the level of capsidiol was increased dramatically in young leaves after fungal inoculation, from not detectable in mock control to 50.68 ± 3.10 µg/g fresh leaves at 3 days post inoculation. Capsidiol-reduced or capsidiol-depleted plants, which were generated by silencing *EAH*s or *EAS*s by virus-induced gene silencing, were more susceptible to the fungus. In addition, this sesquiterpene exhibited strong anti-fungal activities against *A. alternata in vitro* when purified from infected plants and applied to fungal growth. Furthermore, an ERF2-like transcription factor was found to positively regulate capsidiol production and plant resistance through the direct transactivation of a capsidiol biosynthetic gene *EAS12*. Taken together, our results demonstrate that capsidiol, a phytoalexin highly accumulated in *N. attenuata* plants in response to *A. alternata* infection, plays an important role in pathogen resistance independent of JA and ethylene signaling pathways, and its biosynthesis is transcriptionally regulated by an ERF2-like transcription factor.

**Highlight:** Our results demonstrate that capsidiol, a phytoalexin highly accumulated in *Nicotiana attenuata* plants in response to *Alternaria alternata* attack, plays an important role in pathogen resistance, and its biosynthesis is transcriptionally regulated by an ERF2-like transcription factor.

## Introduction

Plants are constantly attacked by a wide variety of microbial pathogens. In response, they activate a large number of intricate defense mechanisms, including the formation of reactive oxygen species, physical reinforcement of cell walls, production of phytohormones, antimicrobial proteins and metabolites (Glazebrook, 2005; Ahuja *et al*., 2012; Mengiste, 2012). The class of small molecules known as phytoalexins is produced by plants *de novo* in response to pathogen attack, and is an important part of the plant defense repertoire (Ahuja *et al*., 2012). In *Arabidopsis*, mutants impaired in the production of the phytoalexin camalexin are more susceptible to infection by necrotrophic fungi, such as *Alternaria brassicicola* (Nafisi *et al*., 2007), *Botrytis cinerea* (Kliebenstein *et al*., 2005) and *Pletosphaerella cucumerina* (Sanchez-Vallet *et al*., 2010). In *Nicotiana* species, two phytoalexins have recently received attention: scopoletin (El Oirdi *et al*., 2010; Sun *et al*., 2014b) and capsidiol (Mialoundama *et al*., 2009; Shibata *et al*., 2010; Grosskinsky *et al*., 2011; Shibata *et al*., 2016).

Capsidiol has been proposed to be an important ‘chemical weapon’ against pathogens in *Nicotiana* species. This bicyclic sesquiterpene is produced via cyclization of farnesyl pyrophosphate (FPP) to 5-*epi*-aristolochene by 5-*epi*-aristolochene synthase (EAS), followed by two hydroxylation reactions catalyzed by 5-*epi*-aristolochene dihydroxylase (EAH) (Facchini and Chappell, 1992; Ralston *et al*., 2001). Capsidiol exhibits toxicity towards many pathogens, including *Phytophthora capsici* and *B. cinerea* (Stoessl *et al*., 1972; Ward *et al*., 1974). Recently, molecular evidence also supports its role in non-host resistance against *P. infestans* in *N. benthamiana*, as *NbEAS*- or *NbEAH*-silenced plants were highly susceptible (Shibata *et al*., 2010).

Ethylene response factors (ERFs) are transcription factors that play crucial roles in plant immunity (Huang *et al*., 2016). In *Arabidopsis*, several ERFs have been identified as important regulators in *Botrytis* resistance, such as ORA59, ERF1 and RAP2.2 (Solano *et al*., 1998; Berrocal-Lobo *et al*., 2002; Pre *et al*., 2008; Zhao *et al*., 2012). Most ERFs are able to bind specifically to DNA sequences containing a GCC (GCC box) and/or a dehydration-responsive element/C-repeat (DRE/CRT box, A/GCCGAC) (Hao *et al*., 1998; Hao *et al*., 2002). However, the ERF RAP2.2 binds the consensus sequence ATCTA in the promoter region of *phytoene synthase* and *phytoene desaturase* to regulate carotenoid biosynthesis (Welsch *et al*., 2007).

The necrotrophic fungal pathogen *Alternaria alternata* (tobacco pathotype) causes brown spot disease in *Nicotiana tabacum* (LaMondia, 2001) and many other *Nicotiana* species, including the wild tobacco *Nicotiana attenuata* (Schuck *et al*., 2014). Inoculation with *A. alternata* elicits the activation of both jasmonate and ethylene signaling pathways in *N. attenuata* plants, which subsequently lead to the accumulation of the phytoalexins, scopoletin and its glycoside form, scopolin (Sun *et al*., 2014b; Li and Wu, 2016; Sun *et al*., 2017). Currently it is not known if fungus-inoculated *N. attenuata* plants produce other phytoalexins, such as capsidiol. If yes, how is its biosynthesis transcriptionally regulated?

In this study, capsidiol was identified and confirmed to be an important phytoalexin produced in *N. attenuata* when challenged by *A. alternata*, and its regulation by an ERF2-like transcription factor was investigated in detail.

## Materials and Methods

### Plant and fungal materials

Seeds of the 31^st^ generation of an inbred line of *Nicotiana attenuata* were used as the wild-type (WT) genotype. Ethylene deficient and insensitive (irACO and Ov-etr1, respectively), and jasmonate deficient (irAOC) *N. attenuata* plants were generated previously (von Dahl *et al*., 2007; Kallenbach *et al*., 2012). Seed germination and plant growth were conducted as described by Krügel *et al*., (2002).

*Alternaria alternata* were grown and inoculated into leaves as described by Sun *et al*., (2014a).

### RNA-seq data processing and analysis

Source-sink transition leaves of rosette-staged WT plants (35-day-old) were detached and inoculated with *A. alternata* for 1 d, when only a few of fungal hyphae had penetrated into leaf tissues (Sun *et al*., 2014a). Total RNA of 3 biological replicates mock (WT_0L_M, with sample names S716, S717 and S719) or inoculated leaf samples (WT_0L_Inf, with sample names S20, S722 and S726) were isolated with TRIzol reagent (Invitrogen). RNA sequencing was conducted by Shanghai OE-Biotech (http://www.oebiotech.com/) with Illumina Hiseq 2000.

Sequencing was performed at 8 G depth, and mapped to the *N. attenuata* reference genome sequence. The relative abundance of the transcripts was measured with the FPKM (RPKM) method, which measures the transcripts abundance as RPKM (Reads Per Kilobase of exon model per Million mapped reads). The differential expressions between mock and inoculated 1 dpi samples with a cutoff of two-fold change, and its significance, were calculated.

### Generation of *NaEAS*- and *NaEAH*-silenced VIGS plants

A 579 bp *NaEAS* cDNA fragment amplified with primers (Z003_F and Z004_R, Supplementary Table S3) and a 424 bp fragment of *NaEAH* cloned with primers (Z047_F and Z048_R, Supplementary Table S3) were individually inserted into pTV00 (Ratcliff *et al*. 2001) in reverse orientations. *Agrobacterium tumefaciens* (strain GV3101) harboring these constructs were mixed with the strain with pBINTRA and inoculated into *N. attenuata* leaves generating *NaEAS*s- and *NaEAH*s-silenced plants (VIGS NaEASs and VIGS NaEAHs). The *A. tumefaciens*–mediated transformation procedure was performed as described previously (Saedler and Baldwin, 2004). To monitor the progress of VIGS, *phytoene desaturase* (PDS) was also silenced. Silencing *PDS* results in the visible bleaching of green tissues (Saedler and Baldwin, 2004; Wu *et al*., 2008) about 2 to 3 weeks after the inoculation. When the leaves of *PDS*-silenced plants began to bleach, the young leaves of VIGS plants and empty vector-inoculated plants (EV plants) were selected for further experiments. Around 20 plants were inoculated with each construct, and usually 10 biological replicates per construct exhibiting efficient silencing were used for each experiment, and all VIGS experiments were repeated twice.

### Purification and quantification of capsidiol in *N. attenuata* after infection

Around 500 g leaves which had been inoculated with *A. alternata* for 3 d, were collected for capsidiol extraction. Leaves were twice extracted with 70 % acetone (2 L) at room temperature. The solvent was evaporated and suspended in water, and then extracted with ethyl acetate. The ethyl acetate-soluble fraction (10 g) was decolorized on MCI gel (www.gls.co.jp) with methanol: H_2_O (90:10) to obtain a yellow gum (7 g), which was subsequently purified by silica gel column with a chloroform: acetone gradient system (from 10:0, 9:1, 8:2, 7:3, 6:4, to 1:1) to yield six main fractions (A**–**F). Fraction B (chloroform: acetone, 9:1; 2 g) was subjected to repeated chromatography over silica gel (petroleum ether: acetone, from 30:1 to 1:1) to yield fractions B1-B4. Fraction B3 (petroleum ether: acetone, 10:1) was separated further by RP-18 column (acetonitrile: H_2_O, 30:70). The obtained crude capsidiol was further purified by semi-preparative HPLC (3 mL/min, UV detection at λ_max_ = 202 nm, acetonitrile: H_2_O, 40:60; ZORBAX SB-C18 column (5 μm, 9.4 × 250 mm, Agilent 1200, USA) to yield capsidiol (30 mg, > 99.5 % purity). The purified capsidiol showed the same characteristic NMR data and HPLC retention times as compared with the authentic standard provided by Prof. Joe Chappell (University of Kentucky, USA).

*A. alternata*-elicited capsidiol levels were determined by HPLC by reference to the authentic capsidiol standard. Each leaf was inoculated with four agar plugs with fungal mycelium, and a 1.5 × 1.5 cm^2^ area of the leaf lamina was harvested at 1 or 3 dpi (around 200 mg fresh mass). Samples were grounded in liquid nitrogen and twice extracted with 1 mL dichloromethane. 2 mL extracts were dried in a SpeedVac concentrator (Eppendorf), and finally dissolved in 1 mL methanol for HPLC. At a flow rate of 1 mL/min, 10 μL of each sample was injected onto a ZORBAX SB-C18 column (5 μm, 4.6 × 250 mm) (Agilent 1260, USA). The mobile phase was composed of solvent A (water) and solvent B (acetonitrile) was used in a isocratic elution (40% of B). Capsidiol was detected at 202 nm, with a retention time of 10.38 min. The standard capsidiol was dissolved in methanol at six concentrations (6.25, 12.5, 25, 50, 100, and 200 *μ*g/mL) to create an external standard curve which was used to calculated fungal-induced capsidiol levels.

### Bioassays for the inhibition of *A. alternata* growth by capsidiol *in vitro*

The inhibition of *A. alternata* mycelium growth by capsidiol *in vitro* was tested in Petri dishes by sub-culturing a 3 mm diameter mycelium plug on PDA medium containing various concentrations of capsidiol for 6 days in the dark at 25□. Twenty mg capsidiol was dissolved in 10 mL methanol, and added to the PDA media at final concentrations of 0, 50, 100 and 200 µg/mL. PDA plates with 1% methanol were served as controls. Photos were taken every two days, and the area of mycelium growth was calculated with ImageJ (http://imagej.nih.gov/ij/).

### Quantification of scopoletin and scopolin

Around 0.2 g leaf samples at 6 dpi of EV, VIGS NaEAS and VIGS NaEAH plants were harvested and ground to fine powder in liquid nitrogen. The levels of scopoletin and scopolin were determined by HPLC-MS/MS as described in Sun *et al*., (2014b).

### Real-Time PCR

Total RNA was extracted from a 1.5 × 1.5 cm^2^ area of leaf lamina which encompassed the inoculation site with TRI reagent (Invitrogen). For each treatment, 4-5 replicate biological samples were collected. cDNA was synthesized from 500 ng total RNA with reverse transcriptases (Thermo Scientific). Real-time PCR was performed on a CFX Connect qPCR System (Bio-Rad) with iTaq Universal SYBR Green Supermix (Bio-Rad) and gene-specific primers as described (Wu *et al*., 2013). For each analysis, a linear standard curve (obtained from threshold cycle number versus log DNA quantity) was constructed by using a dilution series of a specific cDNA sample, and the transcript levels of unknown samples were calculated according to the standard curve. Finally, the relative transcript levels of target genes were obtained by dividing the extrapolated transcript levels of the target genes by the levels of a housekeeping gene *NaActin 2* as an internal standard from the same sample. The transcript abundance of *NaActin 2* was not altered in leaves inoculated with *A. alternata* at 1 and 3 dpi (Xu *et al*., 2018). All primers were listed in Supplementary Table S3.

### Transient expression assays

To determine the subcellular localization of NaERF2-like, a 35S::ERF2-eGFP construct was used for transient expression in *N. attenuata* protoplasts. The Full-length coding sequence of *NaERF2-like* was amplified by primers Z141_F and Z142_R (Supplementary Table S3) and inserted into vector pM999 via *Sac* I and *Xho* I. The method of protoplast isolation and transient transformation was adopted from (Yoo *et al*., 2007) with some modifications. In brief, mesophyll protoplasts were isolated from source-sink transition leaves, and twenty microgram of plasmids was transfected into 2×10^5^ protoplasts with polyethylene glycol (PEG) solution (0.4 g/ml PEG 4000, 0.2 M mannitol, 0.1M CaCl_2_). Transformed cells were cultured in solution (4 mM MES, 0.5 M mannitol, 20 mM KCl) for 18 h in dark and images were obtained with a fluorescent microscope (Leica DM5500 B) with excitation at 488 nm for GFP signal.

For transient transactivation assay, the promoter region of *NaEAS12* (−1671 to −1; numbered from the first ATG) and firefly luciferase (LUC) were amplified and cloned into pCAMBIA1301. Next, the PCR fragment including promoter region of *NaEAS12, LUC*, and Nos terminator was subcloned into pMD18 via *Hind* III and *Sac* I to reduce the size of the vector and increase the transformation efficiency. Similarly, 35S:: NaERF2-like-2HA-eGFP with Nos terminator was also subcloned into pMD18 vector. *N. attenuata* protoplasts were prepared as describe above, and transformed with both NaEAS12_Promoter_::LUC and 35S:: NaERF2-like-2HA-eGFP, or with NaEAS12_Promoter_::LUC and 35S:: 2HA-eGFP as control. After transformation, the protoplasts were subjected to RNA extraction with a PrimeScript™ RT reagent Kit with gDNA Eraser and then gene expression assay. All the experiments were repeated twice with similar results.

### Yeast-one-hybrid assay

The Matchmaker yeast-one-hybrid system (Clontech) was used to test the binding of NaERF2-like and the NaEAS12 promoter *in vitro* according to the user manual. The promoter region of *NaEAS12* (pEAS12-b; −926 to −699; numbered from the first ATG) which contained the candidate binding motif of EM13 (5’-tagattATCTaattctact-3’), was inserted into pAbAi vector with *Hind* III and *Xho* I. The bait construct was linearized and integrated into the genome of yeast strain Y1HGold. The full-length coding sequence of *NaERF2-like* was introduced into pGADT7 AD vector via *Cla* I and *Xho* I, and then the construct was transformed into the yeast cells containing the bait. The positive clones were analyzed on SD/-His/-Leu medium supplied with 200 ng/mL (final conc.) 3-amino-1, 2, 4-trazole (3-AT). Y1HGold [pGADT7/ pEAS12-b-AbAi] was used as negative control, and Y1HGold [pGADT7 Rec-p53/p53-AbAi] was used as positive control.

### Electromobillity shift assays (EMSA)

The full-length coding sequence of *NaERF2-like* was cloned in frame into the *EcoR* I-*Xho* I sites of the pET28a (+), His-NaERF2-like were expressed and purified with Ni-NTA agarose (QIAGEN). Biotin labeled probe EM13 (5’-tagattATCTaattctact-3’) and mutant probe (5’-tagattAATTaattctact-3’) were synthesized from Sangon Biotech (Shanghai). The detection of the binding of the recombinant protein and the probes (300 ng of recombinant protein and 30 ng labeled probe) were carried out with a chemiluminescent EMSA kit (Beyotime Biotechnology) according to the protocol suggested by the manufacture.

### Generation of Ov-NaERF2-like plants and chromatin immune-precipitation assay

The full-length cDNA of NaERF2-like with two HA flags was cloned into pCAMBIA1301 vector after 35S promoter via in-fusion technique (Clontech). *N. attenuata* plants (31^st^ generation of an inbred line) were transformed by *Agrobacterium tumefaciens* with this construct according to Krügel *et al*., (2002). T1 seeds were screened for single T-DNA inserts (1:3 segregation of hygromycin resistance), and two lines of T2 (Ov-NaERF2-like line 1 and 2) with HA signals (Supplementary Fig. S2) were selected and used in this study.

Chromatin immunoprecipitation was performed with EpiQuik Plant ChIP Kit (EPIGENTEK) according to it user manual. The source-sink transition leaves of Ov-NaERF2-like line 2 at 1 dpi (1 g) were used for ChIP assays. A ChIP grade anti-HA antibody (Abcam) was used to immunoprecipitate the protein-DNA complex, and the precipitated DNA was further purified for real time PCR by using primer sets from *NaEAS12* promoter (5^’^-CACTTTAACCCCCGGGTAACT-3^’^ and 5^’^-CACTTCTCAGATTCTCCAGTTTGG-3^’^) and *NaActin* 2 gene (5’-GGTCGTACCACCGGTATTGTG-3’ and 5’-GTCAAGACGGAGAATGGCATG-3’) for negative control. The ChIP experiments were performed twice with similar results. Chromatins which were precipitated without antibody served as the negative controls.

## Results

### Transcriptome analysis reveals the strong regulation of sesquiterpene biosynthetic genes in *N. attenuata* after *A. alternata* inoculation

Previously, we have demonstrated that, in response to *A. alternata* inoculation, *N. attenuata* plants activate both jasmonate and ethylene signaling pathways to regulate the biosynthesis of the phytoalexins scopoletin and its β-glycoside form, scopolin (Sun *et al*., 2014b; Li and Wu, 2016; Sun *et al*., 2017). As *feruloyl-CoA 6’-hydroxylase 1* (*NaF6’H1*), the key enzyme gene for scopoletin biosynthesis, was highly elicited at 1 day post inoculation (1 dpi) when the infection was at early stage and only a few of fungal hyphae had been observed to penetrate into leaf tissues via stomata (Sun *et al*., 2014a), transcriptome analysis was performed in *A. alternata*-inoculated *N. attenuata* leaves at 1 dpi to identify secondary metabolites which could potentially act similarly to scopoletin. Notably, a set of genes involved in terpene synthesis were strongly up-regulated, including *thiolase, HMG-CoA synthase, HMG-CoA reductase, mevalonate kinase, phosphomevalonate kinase, diphosphomevalonate decarboxylase* (*MVPP decarboxylase*), *isopentenyl-diphosphate delta-isomerase, FPP synthase*, the capsidiol biosynthetic genes *5-epi-aristolochene synthases* (*EAS*s) and *5-epi-aristolochene 1,3-dihydroxylases* (*EAH*s), and the solavetivone biosynthetic genes *premnaspirodiene oxygenase* and *premnaspirodiene synthase* (Fig. 1 and Supplementary Table S1). Meanwhile the *squalene synthase* involved in triterpene biosynthesis was down-regulated (Fig. 1 and Supplementary Table S1). These results strongly indicate that sesquiterpene biosynthetic pathway is activated during the inoculation of *A. alternata*.

**Fig. 1.**
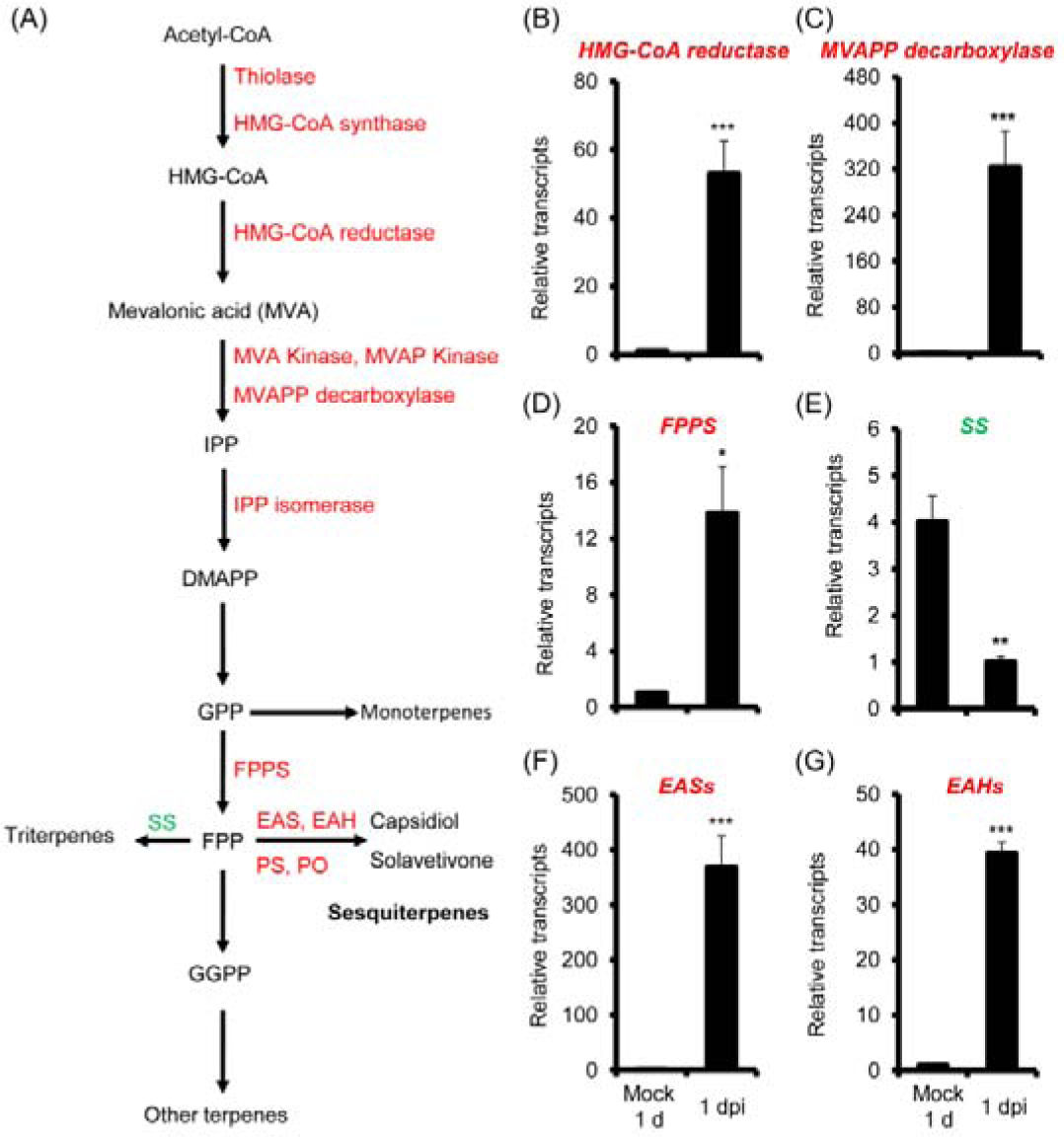
The regulation of terpene biosynthetic genes in *N. attenuata* in response to *A. alternata* inoculation at 1 day post inoculation (dpi) Genes involved in terpene synthesis were strongly regulated during transcriptome analysis in 3 biological replicate samples of mock and 1 dpi: transcripts of all enzymes marked in red font were up-regulated at 1 dpi, while *squalene synthase* (*SS*) with blue font was down-regulated (A). To validate this regulation, relative mean transcripts (± SE) of *HMG-CoA reductase* (A), *MVAPP decarboxylase* (B), *FPPS* (C), *squalene synthase* (D), *EAS*s (E) and *EAH*s (F) were measured by real-time PCR in 4 biological replicate 0 leaves at 1 dpi. Leaves without inoculation were collected as controls (Mock 1 d). Both *EAS*s and *EAH*s were detected by primers conserved in gene family members. Asterisks indicate the level of significant differences between Mock 1 d and 1 dpi samples (Student’s *t*-test: *, *p*<0.05; **, *p*<0.01; ***, *p*<0.005). Enzymes/substrates: 3-hydroxy-3-methyl-glutaryl-CoA (HMG-CoA), 3-hydroxy-3-methyl-glutaryl-CoA synthase (HMG-CoA synthase), 3-hydroxy-3-methyl-glutaryl-CoA reductase (HMG-CoA reductase), mevalonic acid kinase (MVA kinase), mevalonic acid 5-phosphate kinase (MVAP kinase), mevalonic acid 5-diphosphate decarboxylase (MVAPP decarboxylase), isopentenyl diphosphate (IPP), 3’3-dimethylallyl diphosphate (DMAPP), geranyl diphosphate (GPP), farnesyl pyrophosphate synthase (FPPS), farnesyl diphosphate (FPP), 5-*epi*-aristolochene (EA), 5-*epi*-aristolochene synthase (EAS), 5-*epi*-aristolochene hydroxylase (EAH), premnaspirodiene synthase (PS), premnaspirodiene oxidase (PO), geranylgeranyl diphosphate (GGPP).

To confirm the regulation of terpene biosynthesis genes in response to *A. alternata*, we performed quantitative real time PCR on *N. attenuata* samples collected at 1 dpi. Compared with mock infection controls, the transcripts of *MVPP decarboxylase, HMG-CoA reductase, FPP synthase* at 1 dpi were increased to 322-, 53-, and 14-fold respectively, while the transcripts of *squalene synthase* were reduced by 75% (Fig. 1). More importantly, transcripts of the capsidiol biosynthetic genes *NaEAS*s and *NaEAH*s, both of them were encoded by a multi-gene family (Supplementary Table S1) and thus were detected by primers designed at the consensus region, increased to 368- and 40-fold compared to control (Fig. 1). These data fully confirmed the transcriptomics result of strong activation of the sesquiterpene biosynthetic pathway in *N. attenuata* after inoculation.

### Accumulation of capsidiol in *N. attenuata* in response to *A. alternata* inoculation

Since *NaEAS* and *NaEAH* are key genes in the capsidiol biosynthesis pathway, we expected to see an increase in capsidiol production after *A. alternata* inoculation. The levels of capsidiol at 1 and 3 dpi in the young source-sink transition leaves (0 leaves) and mature leaves (+3 leaves) were determined by a high-performance liquid chromatography (HPLC). The +3 leaves are three phyllotaxic positions older than 0 leaves, and are more susceptible to *A. alternata* (Sun *et al*., 2014a). Our results indicated that capsidiol was not detectable in the 0 leaves of mock control, but its level was increased to 4.45 ± 0.72 µg/g fresh leaves in 0 leaves at 1 dpi, and to 50.68 ± 3.10 µg/g at 3 dpi (Fig. 2). Interestingly, this compound was also detected in the susceptible +3 leaves, but its level was only about one-third of that of 0 leaves, with 1.66 ± 1.00 µg/g fresh leaves at 1 dpi, and 14.54 ± 4.39 µg/g at 3 dpi (Fig. 2). These results indicate that 1) capsidiol is highly elicited in *N. attenuata* leaves after inoculation; 2) the lower accumulations of capsidiol in mature leaves may account for their susceptibility to the fungus.

**Fig. 2.**
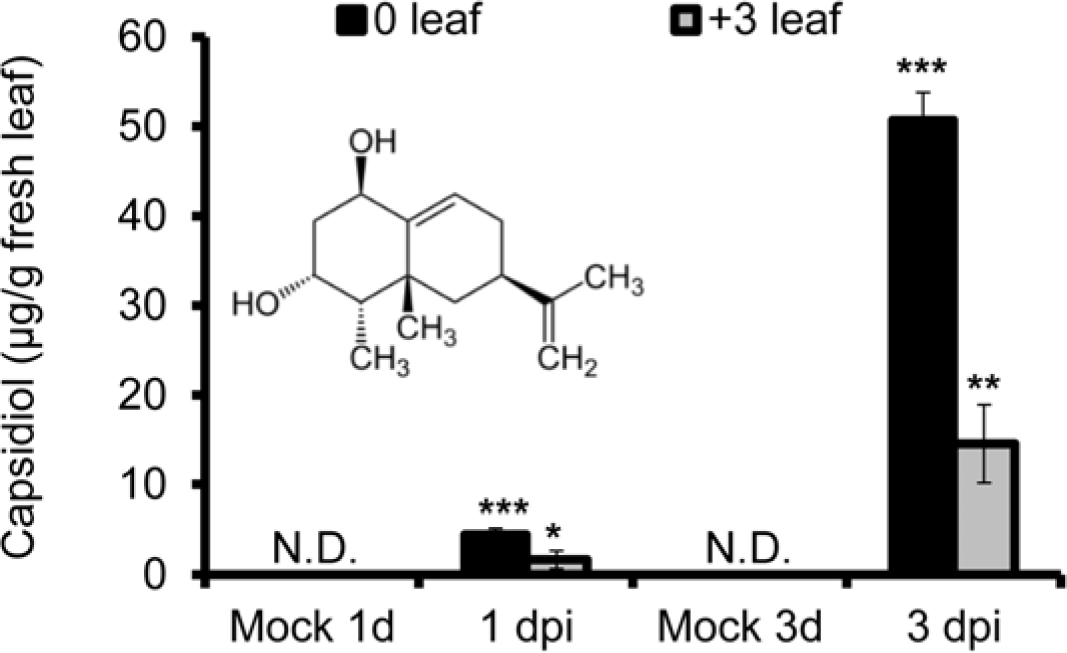
Accumulation of capsidiol in 0 (young) and +3 (mature) leaves after *A. alternata* inoculation. Mean (± SE) capsidiol levels were determined by HPLC in 5 biological replicate 0 and +3 leaves at 1 and 3 dpi. Asterisks indicate levels of significant differences between mock and infected samples (Student’s *t*-test: *, *p*<0.05; **, *p*<0.01; ***, *p*<0.005). N.D., not detectable.

### Capsidiol accumulation is essential for *A. alternata* resistance in *N. attenuata*

To further evaluate the role of capsidiol in *N. attenuata* resistance against *A. alternata*, we silenced *NaEAS*s and *NaEAH*s separately with their conserved sequences, as both are encoded by members of a multi-gene family. Compared with mock controls, transcripts of *NaEAH*s were dramatically induced in young leaves of *N. attenuata* plants transformed with empty vector (EV) at 2 dpi; however, plants transformed with the *NaEAH*s-silencing construct (VIGS NaEAHs) showed a 92% reduction in *NaEAH*s transcripts compared to EV plants with the same treatments, indicating effective silencing of the *NaEAH*s (Fig. 3A). We also investigated capsidiol level in EV and VIGS NaEAHs plants at 3 dpi. Capsidiol levels at 57.72 ± 5.88 µg/g fresh leaves were detected in EV plants, while only 8.99 ± 1.56 µg/g fresh leaves were found in VIGS NaEAHs plants (Fig. 3B).

**Fig. 3.**
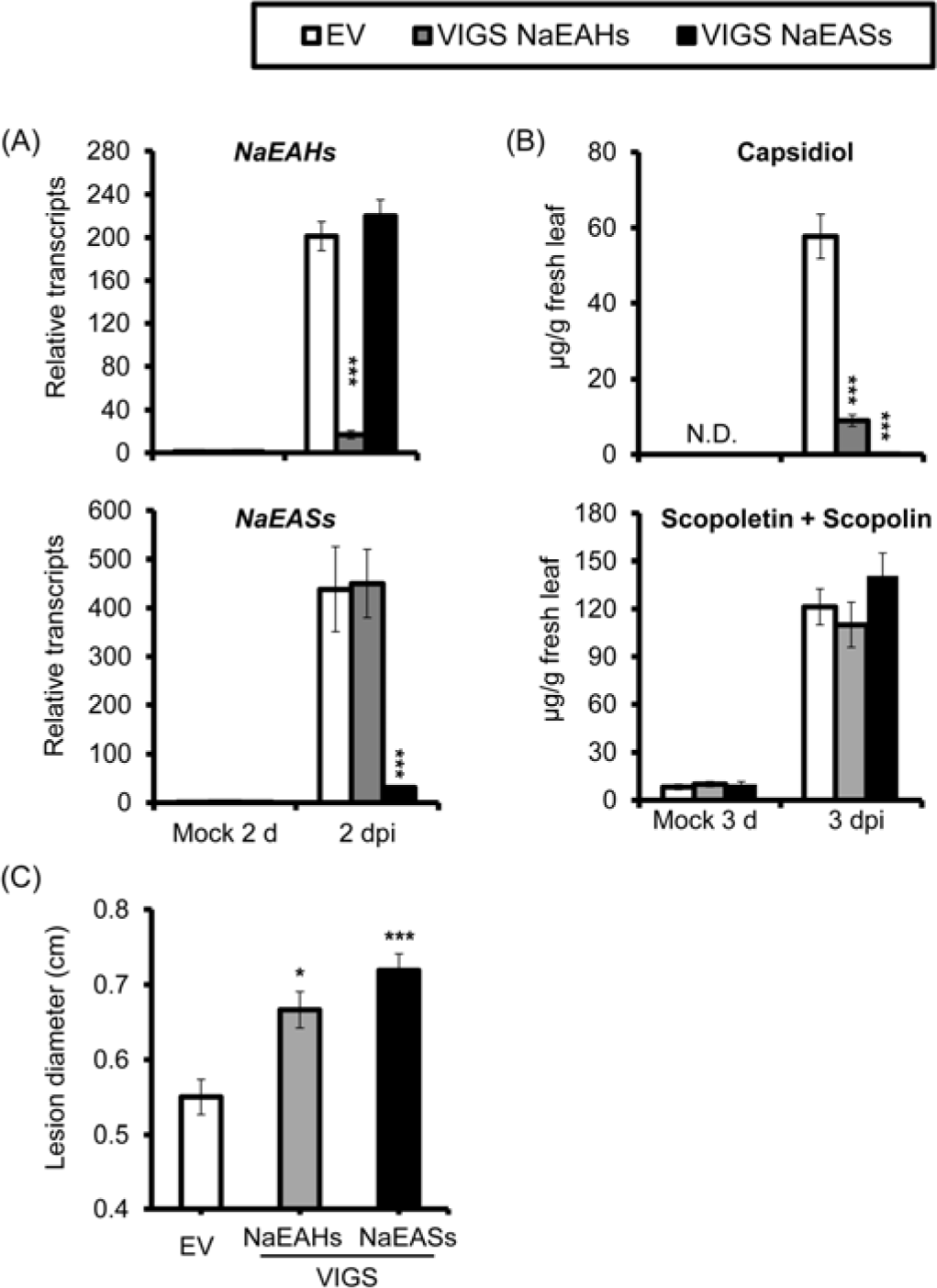
Silencing *NaEAHs* or *NaEAS*s expressions dramatically reduces *A. alternata*-induced capsidiol levels and plant resistance without affecting scopoletin. (A): Mean (± SE) relative *A. alternata*-induced *NaEAH*s and *NaEAS*s transcripts as measured by real-time PCR in 5 replicate young leaves of EV, VIGS NaEAHs and VIGS NaEASs plants at 2 dpi. (B): Mean (± SE) capsidiol and scopoletin (including scopolin) levels were determined by HPLC in 5 replicate young leaves of EV, VIGS NaEAHs and VIGS NaEASs plants at at 3 dpi. (C): Mean (± SE) diameter of necrotic lesions in 8 replicate young leaves of EV, VIGS NaEAHs and VIGS NaEASs plants infected with *A. alternata* for 6 d. As both *NaEAH* and *NaEAS* are encoded by large gene families, conserved cDNA regions were used for silencing and real time PCR. The asterisks indicate levels of significant differences between EV and VIGS leaves (Student’s *t*-test: *, p<0.05; ***, *p*<0.005). N.D., not detectable.

Because a small amount of capsidiol was still present in VIGS NaEAHs plants, we attempted to generate additional capsidiol-depleted *N. attenuata* plants by silencing *NaEAS*s. *NaEAS*s expression was successfully silenced, as only 7% of the transcripts of *NaEAS*s were detected at 2 dpi in VIGS NaEASs plants compared with EV plants (Fig. 3A). More importantly, the *A. alternata*-elicited capsidiol levels at 3 dpi was abolished in VIGS NaEASs plants, with only 0.2 % of the levels detected in EV plants, which were comparable to the levels quantified in the mock controls of EV plants (Fig. 3B). From these data, we infer that *NaEASs* genes are crucial for *A. alternata*-elicited capsidiol production.

To test whether capsidiol-reduced or -depleted plants are more susceptible to *A. alternata*, young leaves of EV, VIGS NaEAHs, and VIGS NaEASs plants were inoculated with the fungus. Two independent VIGS experiments showed significantly increased lesion diameters in VIGS NaEAHs (121% of EV plants) and VIGS NaEASs plants (131% of EV plants) (Fig. 3C). Meanwhile, we did not observe changes in scopoletin and scopolin, two important phytoalexins involved in *A. alternata* resistance (Fig. 3B). These results strongly indicate that capsidiol plays an important role in defending against *A. alternata*.

### Capsidiol exhibits anti-fungal activity against *A. alternata in vitro*

To test whether capsidiol has a direct impact on fungal growth or not, we purified 30 mg capsidiol from 500 g of *A. alternata*-inoculated leaves, and applied this compound in various concentrations to the growth medium to evaluate its inhibition activity of fungal growth *in vitro*. The fungi were grown on PDA plates containing with 50 µg/mL or 100 µg/mL capsidiol, and we observed fungal growth was reduced to 56.6% or 43.8% of that of controls. Application of 200 µg/mL capsidiol resulted in further reduction in growth to 37.1% of control (Fig. 4A, B). These results suggest that capsidiol at a concentration observed *in planta* has a direct impact on the fungal growth *in vitro*.

**Fig. 4.**
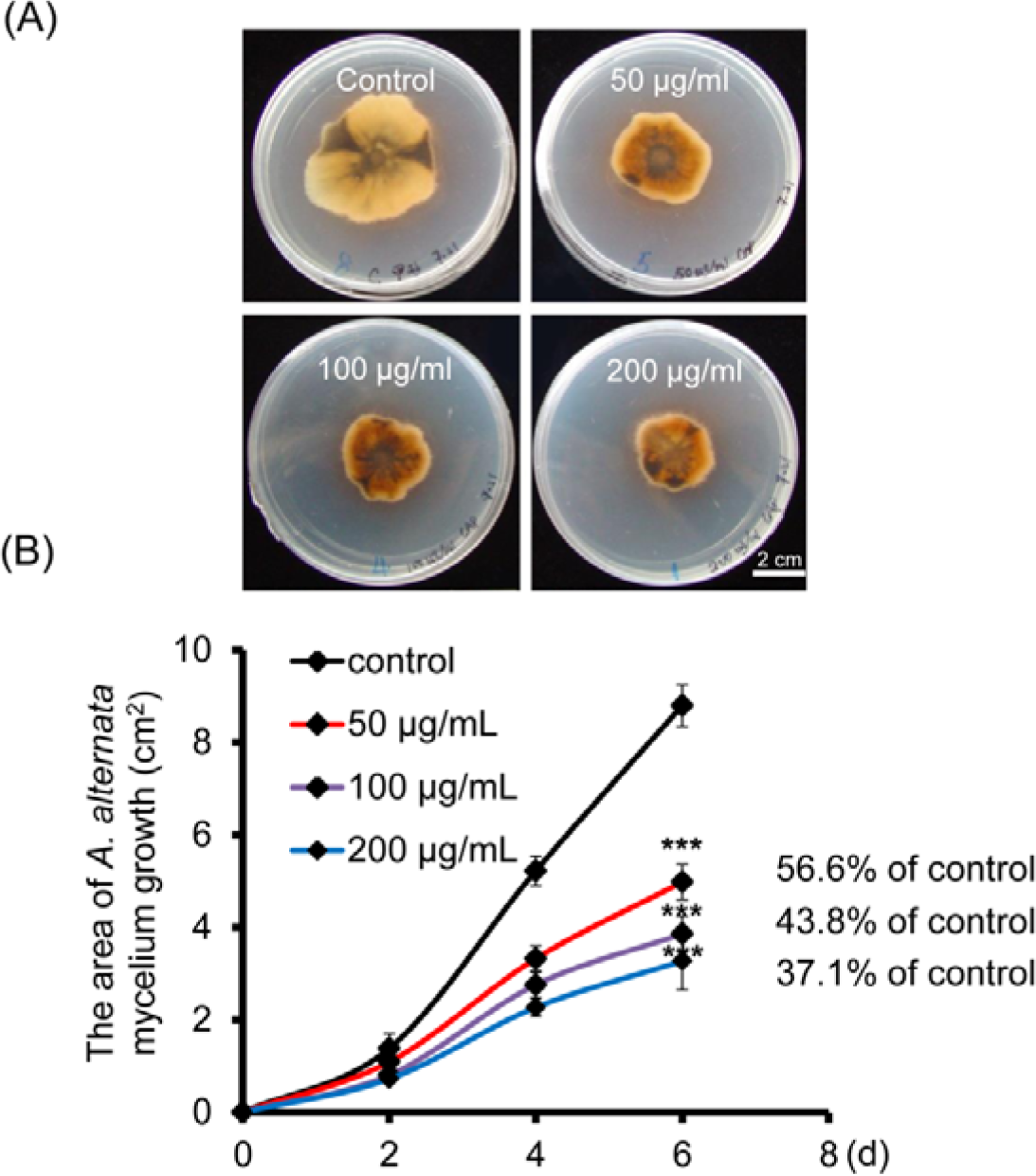
Capsidiol exhibits anti-fungal activity *in vitro*. (A): Growth of *A. alternata* mycelium at day 6 in PDA with capsidiol at final concentration of 0, 50, 100, and 200 µg/mL. PDA plates with 1 % of methanol were served as controls. (B): The area of *A. alternata* mycelium growth in PDA with different concentrations of capsidiol was determined by ImageJ. Data were collected every 2 d. Asterisks indicate levels of significant differences between mock and infected samples (Student’s *t*-test: *, *p*<0.05; **, *p*<0.01; ***, *p*<0.005).

### *A. alternata*-induced capsidiol accumulation is not dependent on JA and ethylene signaling

JA and ethylene signaling pathways are crucial for phytoalexin scopoletin biosynthesis. To investigate the roles of these two signaling pathways in capsidiol biosynthesis, we measured the levels of capsidiol and transcripts of *NaEASs* and *NaEAH*s after *A. alternata* inoculation in WT, JA-deficient (irAOC), ethylene-deficient (irACO), and ethylene-insensitive (Ov-etr1) plants generated previously (Kallenbach *et al*., 2012; von Dahl *et al*., 2007). We found that the induction levels of *NaEASs* and *NaEAHs* by *A. alternata* were similar in WT, irAOC, irACO and Ov-etr1 plants at both 1 and 3 dpi (Fig. 5A, B). In addition, no significant differences of capsidiol production were observed in WT, irAOC, irACO and Ov-etr1 plants at 3 dpi (Fig. 5C). Thus, our data indicated that *A. alternata*-induced capsidiol accumulation is not dependent on JA and ethylene signaling.

**Fig. 5.**
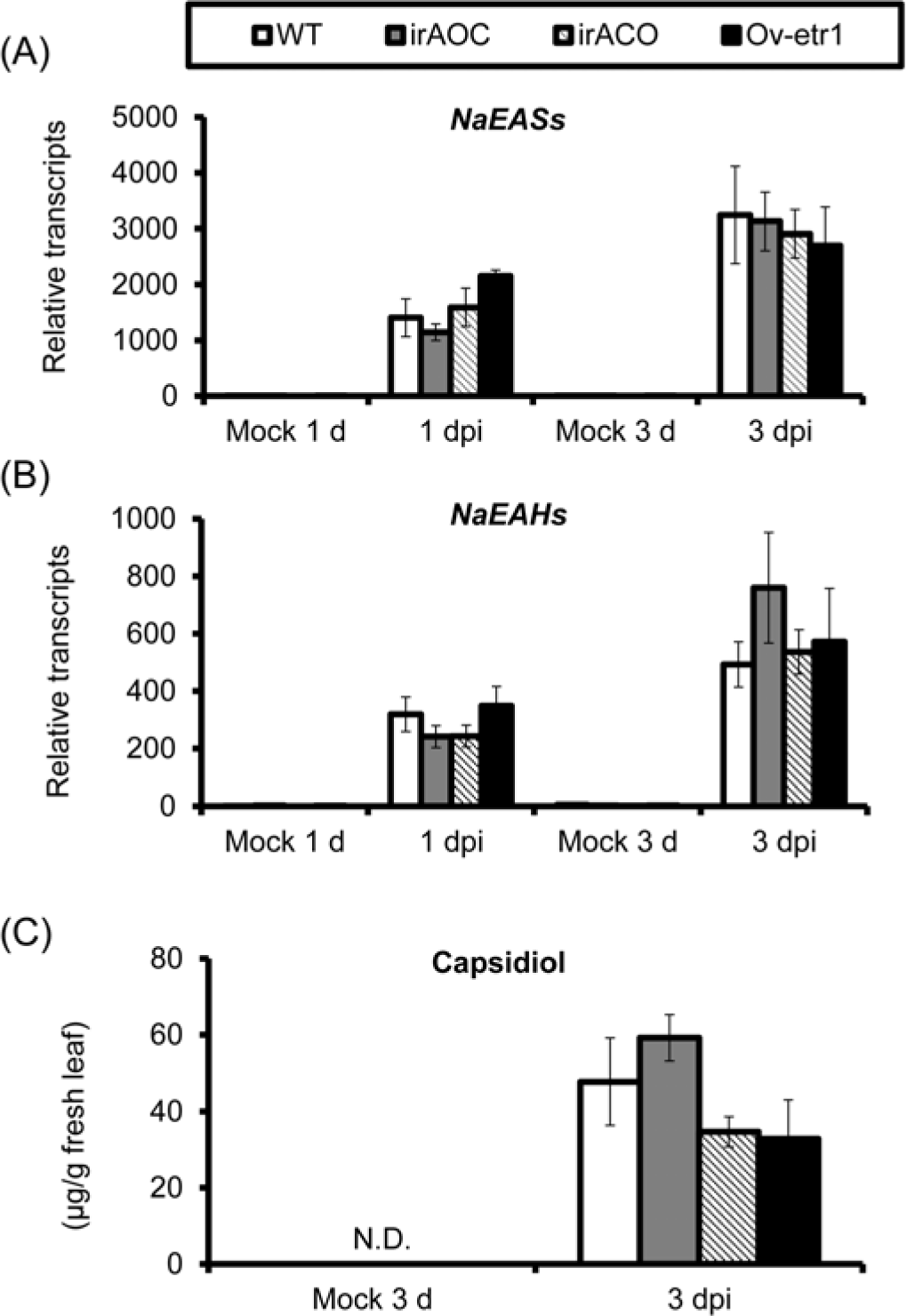
JA and ethylene pathways play a minor role in *A. alternata*-induced transcripts of *NaEAS*s and *NaEAH*s, and capsidiol biosynthesis. Mean (± SE) relative *A. alternata*-induced *NaEAS*s(A)and *NaEAHs*(B)transcripts as measured by real-time PCR in 5 biological replicates of young leaves (0 leaves) in WT, irAOC (JA-deficient), irACO (ethylene deficient) and Ov-etr1 (ethylene-insensitive) plants at 1 and 3 dpi. Mean (± SE) capsidiol levels (C) were determined by HPLC in 5 replicated 0 leaves of WT, irAOC, irACO and Ov-etr1 plants at 3 dpi. N.D., not detectable.

### NaERF2-like, a transcription factor highly induced in young leaves, is required to mount a capsidiol-based defense

Since we observed increases in both capsidiol production as well as *NaEASs* and *NaEAH*s transcripts in *N. attenuata* leaves after fungal inoculation, we hypothesized that the transcription factors regulating *NaEASs* and *NaEAH*s expression were also increased. Thus, we silenced the expression of the 6 fungus-elicited transcription factor genes with the most abundant transcripts after fungal inoculation (Supplementary Table S2), including *ethylene-responsive transcription factor ABR1-like* (*NaERF ABR1-like*; gene accession number : XM_019374371), *zinc finger protein ZAT12-like* (*NaZAT12*; gene accession number: XM_019368773.1), *probable WRKY transcription factor 40* (*NaWRKY40*; gene accession number: XM_019402562.1), *probable WRKY transcription factor 43* (*NaWRKY43*; gene accession number: XM_019375046.1), *probable WRKY transcription factor 61* (*NaWRKY61*; gene accession number: XM_019371308.1), *ethylene-responsive transcription factor 2-like*(*NaERF2-like*; gene accession number: XM_019399671.1), to identify the regulator(s) responsible for capsidiol biosynthesis. *A. alternata*-elicited *NaEAS12* was not altered in plants silenced with *NaERF ABR1-like, NaZAT12, NaWRKY40, NaWRKY43*, or *NaWRKY61* (Supplementary Fig. S1).

*NaERF2-like* (Gene accession number: XM_019399671.1), one of the top 6 transcription factors strongly up-regulated in response to fungal inoculation in our transcriptome analysis, was highly induced in *N. attenuata* 0 leaves at both 1 and 3 dpi (Fig. 6A; Supplementary Table S2). The NaERF2-like protein exhibited nuclear localization in the protoplast of *N. attenuata* when its eGFP fusion protein was driven by a constitutive 35S promoter (Fig. 6B).

**Fig. 6.**
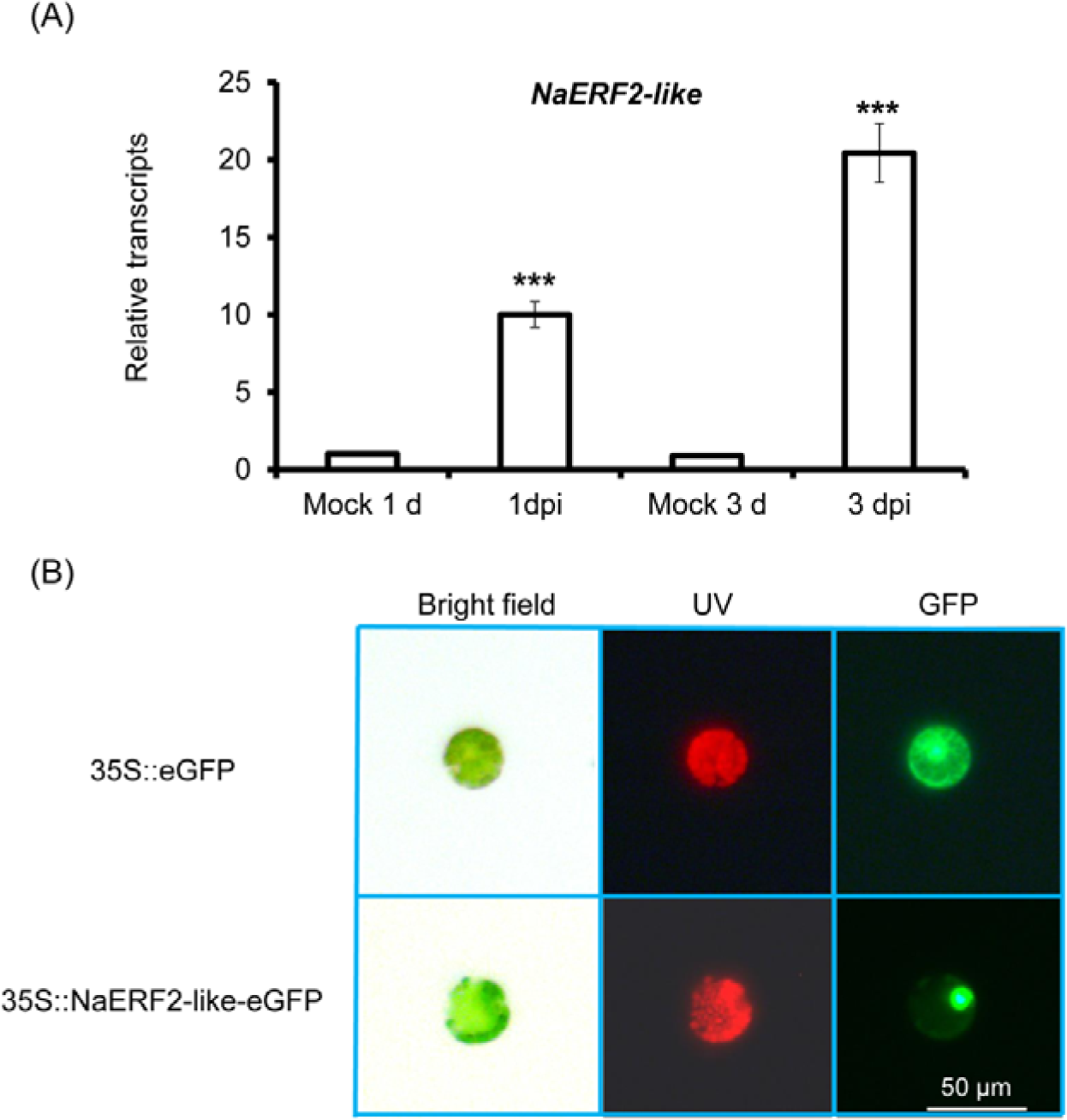
NaERF2-like is highly elicited after *A. alternata* inoculation and is targeted to the nucleus. (A): Mean (± SE) relative *A. alternata*-induced *NaERF2-like* transcripts as measured by real-time PCR in 4 replicate 0 leaves at 1 and 3 dpi. The asterisks indicate levels of significant differences between mock and infected samples (Student’s *t*-test: *, p<0.05;***, *p*<0.005). (B): Two fusion proteins, 35S::eGFP and 35S::NaERF2-like-eGFP, were expressed transiently in *N. attenuata* protoplasts, respectively. NaERF2-like was found to be targeted to the nucleus. Images were taken in bright field (left), UV (middle) and GFP channels (right).

To investigate the role of the *NaERF2-like* transcription factor in capsidiol biosynthesis in detail, we silenced the gene by VIGS, and then measured the levels of capsidiol and transcripts of *NaEASs* and *NaEAH*s after fungal inoculation.

*NaERF2-like* transcripts were highly elicited at 2 dpi in EV plants; in contrast, VIGS NaERF2-like plants showed an 87% reduction in *NaERF2-like* transcripts after the same treatment (Fig. 7A). Compared to EV plants at 2 dpi, the transcripts of *NaEASs* and *NaEAH*s in VIGS NaERF2-like plants were reduced by 75% and 62%, respectively (Fig. 7B, C). As expected, *A. alternata*-induced capsidiol level in VIGS NaERF2-like plants at 3 dpi was reduced by 68% when compared with EV plants (Fig. 7E). However, the fungus-elicited transcriptional levels of *NaF6’H1* were not affected in VIGS NaERF2-like plants (Fig. 7D). These results indicate that silencing *NaERF2-like* substantially decreases *A. alternata*-elicited transcription of *NaEASs* and *NaEAH*s, and consequently, capsidiol level, without affecting scopoletin-based defense.

**Fig. 7.**
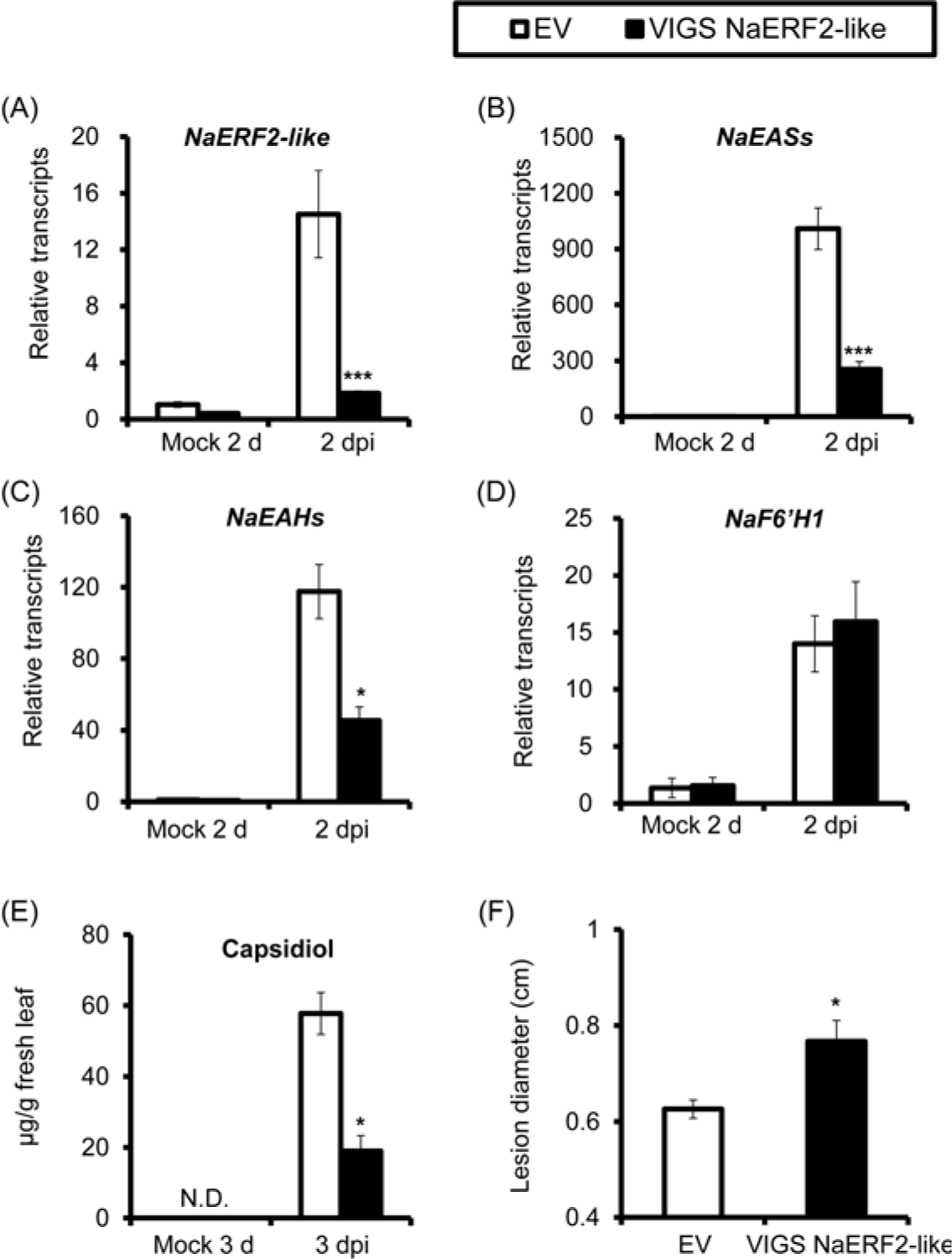
Silencing *NaERF2-like* impairs *A. alternata*-induced transcripts of *NaEAS*s and *NaEAH*s, capsidiol levels and plant resistance without affecting *NaF6’H1* transcript accumulation. Mean (± SE) relative *A. alternata*-induced *NaERF2-like* (A), *NaEAS*s (B), *NaEAH*s and *NaF6’H1* (D) transcript abundance as measured by real-time PCR in 5 replicate young leaves of EV and VIGS NaERF2-like plants at 2 dpi; capsidiol levels (E) were determined by HPLC in 5 replicate young leaves of EV and VIGS NaERF2-like plants at 3 dpi; mean (± SE) diameter of necrotic lesions (F) were recorded in 8 replicate young leaves of EV and VIGS NaERF2-like plants infected with *A. alternata* for 6 d. Two independent VIGS experiments returned similar results. The asterisks indicate levels of significant differences between EV and VIGS leaves (Student’s *t*-test: *, p<0.05; ***, *p*<0.005). N.D., not detectable.

In addition, silencing *NaERF2-like* led to plants more susceptible to *A. alternata*, as significantly larger lesions were observed in VIGS NaERF2-like plants at 6 dpi in two independent VIGS experiments (Fig. 7F).

### NaERF2-like directly regulates the capsidiol biosynthetic gene *NaEAS12*

Next, we explored the mechanism by which NaERF2-like regulates capsidiol-based resistance. We hypothesized that NaERF2-like might directly regulate genes in the capsidiol biosynthetic pathway. Since *NaEAS12* expression was greatly reduced in *NaERF2-like*-silenced plants (Fig. 8A), we selected this gene to test whether its promoter could be directly activated by NaERF2-like.

**Fig. 8.**
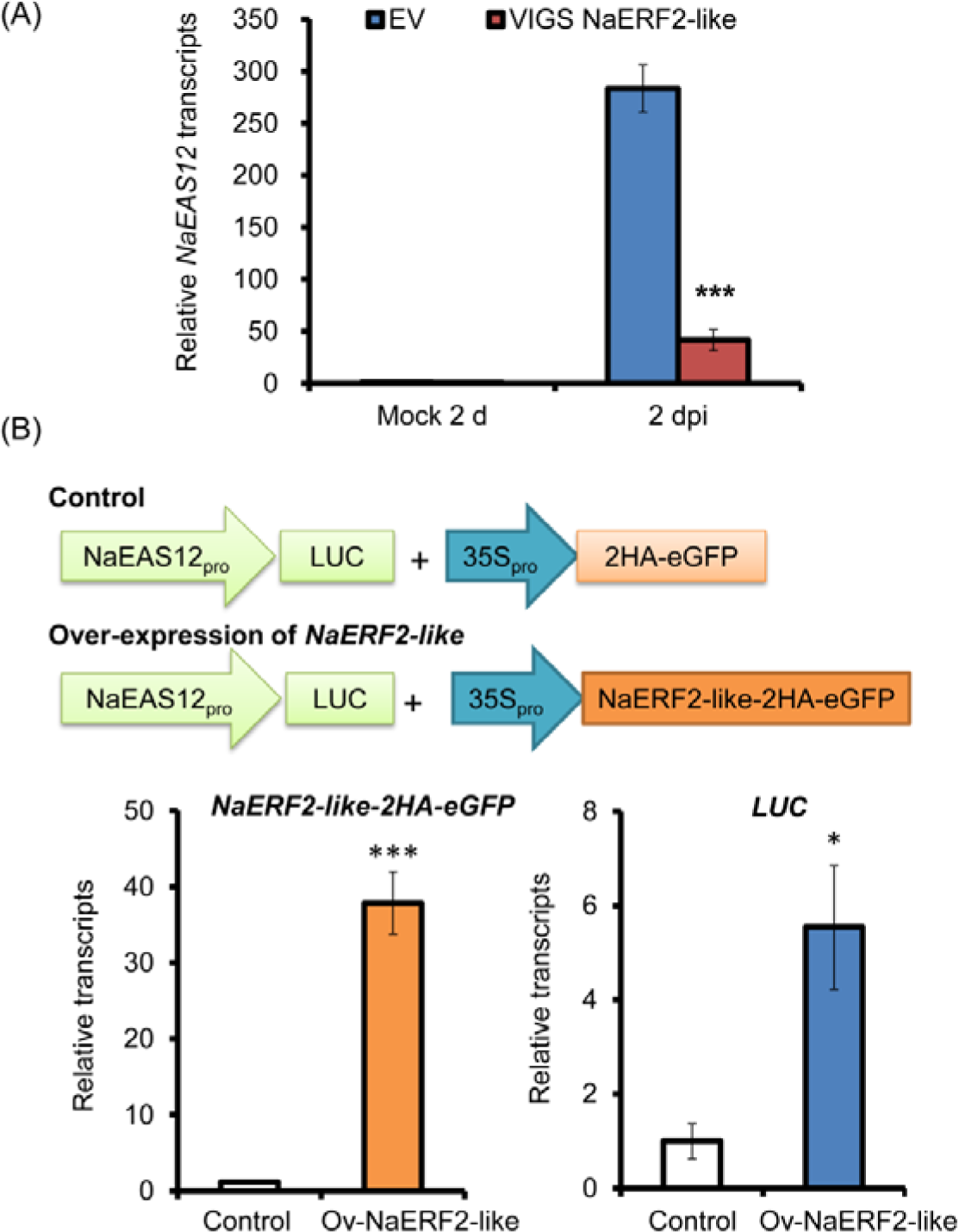
*A. alternata*-elicited *NaEAS12* transcripts are largely dependent on *NaERF2-like*, and transient over-expression of *NaERF2-like* leads to transactivation of *NaEAS12* promoter. (A): Mean (± SE) relative *A. alternata*-induced *NaEAS12* transcripts as measured by real-time PCR in 5 replicate young leaves of EV and VIGS NaERF2-like plants at 2 dpi. The asterisks indicate levels of significant differences between EV and VIGS leaves at 2 dpi (Student’s *t*-test: ***, *p*<0.005). (B): Transcript abundance of *NaERF2-like-2HA-eGFP* and *LUC* in *N. attenuata* protoplasts co-expressing *NaEAS12* promoter::LUC and 35S::2HA-eGFP or 35S:: NaERF2-like-2HA-eGFP, were measured in 3 biological samples. Experiments were repeated twice with similar results. The asterisks indicate levels of significant differences between control and NaERF2-like over-expressing protoplasts (Student’s *t*-test: *, p<0.05; ***, *p*<0.005).

Several lines of evidence were consistent with the idea that NaERF2-like directly regulates the capsidiol biosynthetic gene *NaEAS12*. When the *NaERF2-like* gene was over-expressed in the protoplasts of *N. attenuata, luciferase* gene (LUC) driven by the *NaEAS12* promoter showed a 5.5-fold increase in expression (Figure 8B), indicating that the over-expression of *NaERF2-like* enhanced the transcriptional activity of *NaEAS12* promoter. From the promoter region of *NaEAS12*, an ATCTA motif, previously shown to be the binding site of RAP2.2 in *Arabidopsis* (Welsch *et al*., 2007), was identified and confirmed as the binding site for the ERF2-like by yeast-one-hybrid, electrophoretic mobility shift assay (EMSA) and chromatin immunoprecipitation-based qPCR (ChIP-qPCR). Yeast-one-hybrid experiments revealed that the ERF2-like protein could bind to the *NaEAS12* promoter fragment EAS12-b (located from −699 to −926 bp upstream of the starting codon), which contained the ATCTA motif (Fig. 9). Further EMSA experiments indicated that NaERF2-like could directly bind to the biotin labeled probe EM13 (5′-tagattATCTAattctact-3′), but not to the mutated one (5′-tagattAATTAattctact-3′)(Fig.9). To further confirm these results *in vivo*, we generated a transgenic line ectopically-expressing the NaERF2-like protein fused with two HA tags at the C-terminal (35S:: NaERF2-like-2HA) for use in ChIP-qPCR. The fusion protein could be detected by a commercial HA antibody (Supplemental Fig. S2), suggesting that the protein was successfully over-expressed in *N. attenuata* plants. Transgenic plants were inoculated with *A. alternata* and sampled at 1 dpi. We found that the HA-tagged NaERF2-like protein bound the *NaEAS12* promoter at a site which encompassed the ACTCA motif (Fig. 9).

**Fig. 9.**
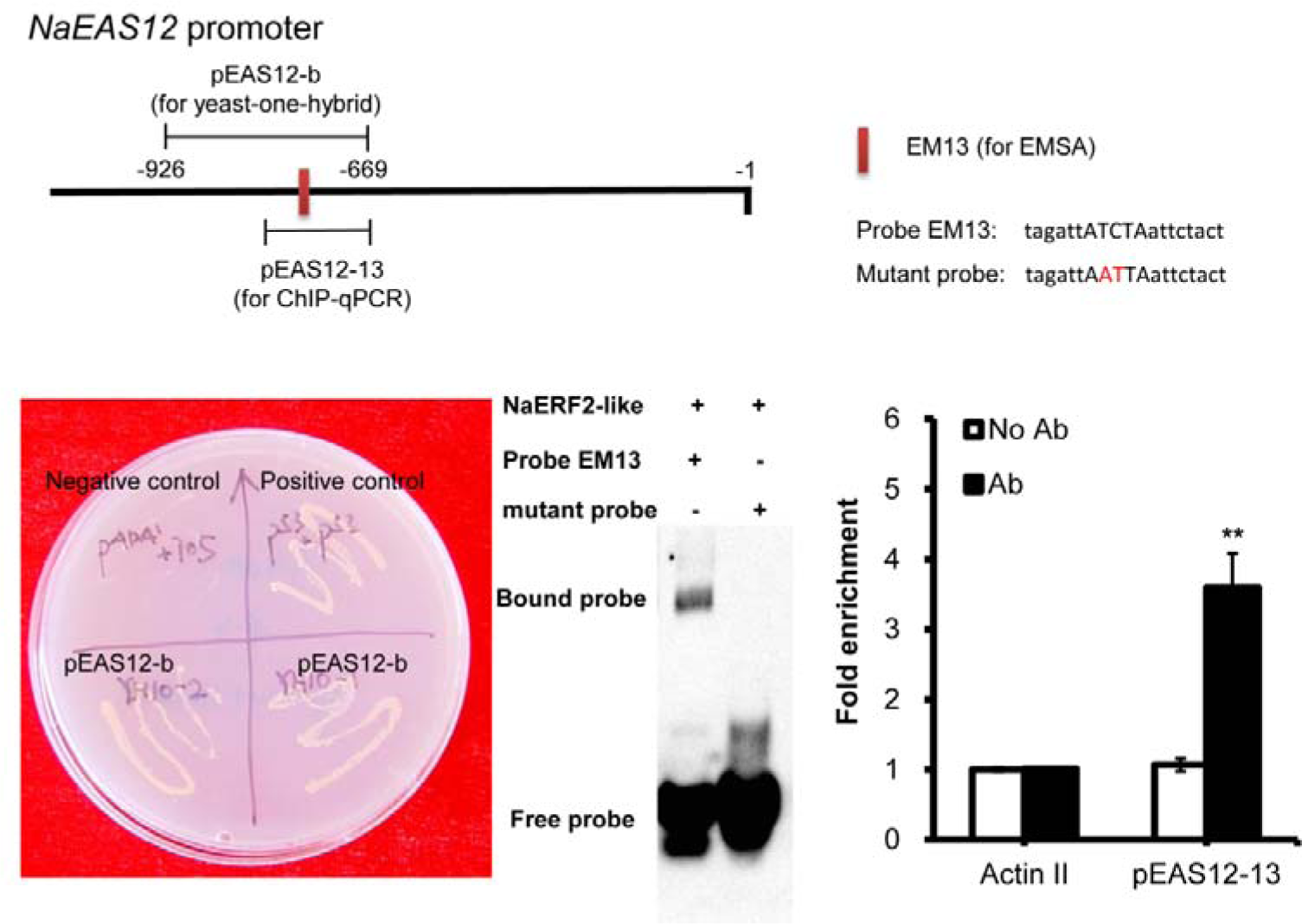
The binding of NaERF2-like and *NaEAS12* promoter as demonstrated by yeast-one-hybrid, EMSA and chromatin immunoprecipitation. The NaEAS12 promoter structure indicated the pEAS12-b (−669 to −926 numbered from the ATG) for yeast-one hybrid, the probe EM13 (5’-tagattATCTaattctact-3’) for EMSA, and the pEAS12-13 (−669 to −781 numbered from the ATG) for ChIP assays. Yeast-one-hybrid analysis revealed that NaERF2-like could bind to EAS12-b as the yeast cells could grow on the SD/-His/-Leu medium supplied with 200 ng/mL (final conc.) 3-AT; Y1HGold [pGADT7/ pEAS12-b-AbAi] was used as a negative control, and Y1HGold [pGADT7 Rec-p53/p53-AbAi] was used as a positive control. EMSA demonstrated that His tagged NaERF2-like could bind to probe EM13 but not to the mutated one. The mutant probe (5’-tagattAATTaattctact-3’) served as a negative control in EMSA. ChIP-real time PCR data indicated NaERF2-like bound to the promoter of *NaEAS12*. Negative controls were without antibody (No Ab) and with HA antibody but using primers detecting *NaActin 2*. The asterisks indicate levels of significant differences between No Ab and with Ab in pEAS12-13 (Student’s *t*-test: **, p<0.01).

### Over-expression of *NaERF2-like* does not alter plant resistance, but increases *A. alternata*-induced *NaEAS12* gene expression and capsidiol levels

To further understand the role of *NaERF2-like* in pathogen defense we investigated *NaEAS12* gene expression, capsidiol level and plant resistance in WT, and two *NaERF2-like* over-expression (Ov-NaERF2-like) lines. The ectopic over-expression of *NaERF2-like* fused with two HA tags substantially increased the basal and induced transcriptional levels of *NaERF2-like* (Fig. 10A) without affecting the plant’s morphology and size. As expected, Ov-NaERF2-like line 1 and Ov-NaERF2-like line 2 showed 178% and 219% of the *NaEAS12* expression of WT when 0 leaves were inoculated at 1 d (Fig. 10A). Consistently, capsidiol levels attained values 149% and 175% of that of WT in Ov-NaERF2-like lines 1 and 2 at 3 dpi, respectively (Fig. 10B). However, we found no difference in lesion diameters in both over-expression lines compared with WT (Supplementary Fig. S2), indicating that over-expression of *NaERF2-like* increased the gene expression of *NaEAS12*, and subsequently capsidiol biosynthesis, but had only a minor effect on the plants’ resistance.

**Fig. 10.**
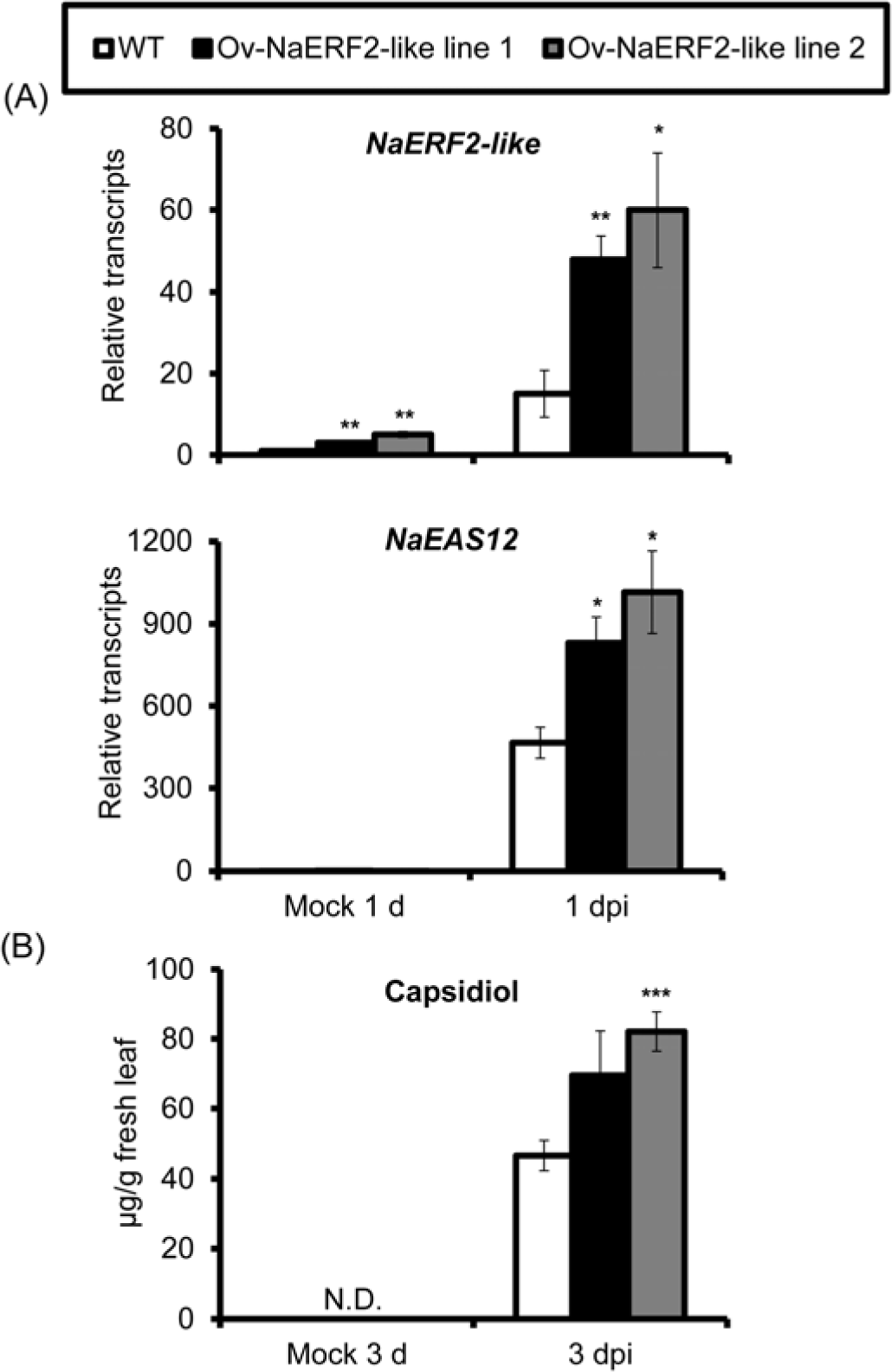
Over-expression of *NaERF2-like* enhances *NaEAS12* expression and capsidiol production in stable transformation plants. (A): Mean (± SE) relative *A. alternata*-induced *NaERF2-like* and *NaEAS12* transcripts as measured by real-time PCR in 5 replicate 0 leaves of WT, Ov-NaERF2-like line 1 and Ov-NaERF2-like line 2 plants at 1 dpi. (B): Mean (± SE) capsidiol levels were determined by HPLC in 5 replicate 0 leaves of WT, Ov-NaERF2-like line 1 and Ov-NaERF2-like line 2 plants at 3 dpi. The asterisks indicate levels of significant differences between WT and Ov-NaERF2-like lines after inoculation (Student’s *t*-test: *, p<0.05; **, p<0.01; ***, *p*<0.005). N.D., not detectable.

## Discussion

Phytoalexins are important ‘chemical weapons’ employed by plants in defending against pathogens. In addition to scopoletin and scopolin, two phytoalexins regulated by JA and ethylene signaling pathways in response to *A. alternata* infection (Sun *et al*., 2014b; Li and Wu, 2016), we demonstrate in this study that capsidiol is another important phytoalexin produced by *N. attenuata*, and its biosynthesis is not dependent on JA and ethylene signaling pathways but is transcriptionally regulated by a transcription factor NaERF2-like.

### Capsidiol is an important phytoalexin produced in *N. attenuata* in response to *A. alternata* infection

Capsidiol was initially isolated from pepper fruit after treatments of various pathogens, including *Phytophthora capsici, Botrytis cinerea*, and *Fusarium oxysporum* (Stoessl *et al*., 1972). Later, this compound was also identified from infected *Nicotiana* species (Bailey *et al*., 1975; Guedes *et al*., 1982; Mialoundama *et al*., 2009; Shibata *et al*., 2010; Grosskinsky *et al*., 2011; Shibata *et al*., 2016). In our *N. attenuata* -*A. alternata* pathosystem, we found that a large number of genes were strongly regulated in response to *A. alternata* inoculation at 1 dpi during transcriptome analysis. Many of these genes were involved in the biosynthesis of sesquiterpenes, capsidiol and solavetivone (Fig. 1 and Supplementary Table S1). Indeed, the levels of capsidiol were dramatically induced to 50.68 ± 3.10 µg/g fresh leaves at 3 dpi in young 0 leaves (Fig. 2). These finding suggest that capsidiol is involved in the resistance of *N. attenuata* to *A. alternata* infection.

Ideally, the benefits of a putative resistant trait should be determined in plants differing only in a single gene that controls the defense trait and are otherwise identical (Bergelson *et al*., 1996). In this study, virus-induced gene silencing of *NaEAH*s or *NaEAS*s was used to manipulate the production of capsidiol, and the results revealed that capsidiol-reduced or -depleted plants were more susceptible to *A. alternata* (Fig. 3). In addition, capsidiol showed strong anti-fungal activities against *A. alternata* when it was extracted and purified from infected plants and applied to fungal growth *in vitro* at a concentration in the range of observed *in planta* (Fig. 4).

Thus, our results demonstrate that capsidiol is an important phytoalexin involved in the defense mechanism of *N. attenuata* against *A. alternata*, and the high level of capsidiol accumulated in the young 0 leaves accounts for their high resistance to *A. alternata* infection. Whether solavetivone plays a role in resistance is unknown. Transcriptome data, especially the strong up-regulation of *premnaspirodiene oxygenases* and *premnaspirodiene synthases* at 1 dpi (Fig. 1 and Supplementary Table S1), is consistent with a defensive role of solavetivone. However, this question needs further investigation.

### Regulation of capsidiol biosynthesis

Several reports indicate that the phytoalexin production is influenced by endogenous plant hormones. Increasing cytokinins (CK) levels in *N. tabacum* plants by exogenous CK application or overexpression of bacterial *isopentenyl transferase* gene enhanced their resistance to the hemibiotrophic bacterium, *Psudomonas syringae*, by increasing levels of capsidiol and scopoletin (Grosskinsky *et al*., 2011). This CK-mediated resistance is independent of salicylic acid, jasmonate and ethylene signaling pathways (Grosskinsky *et al*., 2011). Abscisic acid (ABA) negatively regulates elicitor-induced biosynthesis of capsidiol in *N. plumbaginifolia*; a two-fold increase in capsidiol synthesis was observed in ABA-deficient mutants compared with WT plants when exposed to cellulose or *B. cinerea* (Mialoundama *et al*., 2009). In addition, when *N. benthamiana* was inoculated with *Phytophthora infestans*, pathogen-induced capsidiol and *NbEAS* and *NbEAH* expression were abolished in plants silenced with *ethylene insensitive 2* (*NbEIN2*), suggesting that ethylene signaling pathway is essential for capsidiol production (Shibata *et al*., 2010; Ohtsu *et al*., 2014).

In contrast to the observation in *N. benthamiana* inoculated with *P. infestans*, our experiments did not support the role of ethylene signaling in capsidiol elicitation in *N. attenuata*-*A. alternata* pathosystem. When *NaEIN2* was silenced by VIGS, *A. alternata*-elicited *NaF6’H1* was dramatically reduced, which is consistent with our previous finding that ethylene signaling is essential for scopoletin biosynthesis (Sun *et al*., 2017), but *NaEAH*s and *NaEAS*s trancripts were induced to levels similar to plants transformed with empty vector (Supplementary Fig. S3). This result indicates that blocking the ethylene signaling pathway has little effect on the expression of these two key capsidiol biosynthesis genes. Additional evidence against the involvement of ethylene signaling comes from irACO plants (ethylene-deficient) and Ov-etr1 plants (ethylene-insensitive), both of which were generated previous by von Dahl *et al* (2007). Both capsidiol production and transcripts of *NaEAH*s and *NaEAS*s were induced to the same high levels in WT, irACO and Ov-etr1 plants (Fig. 5). Thus, we concluded that ethylene signaling pathway does not play a critical role in capsidiol production and gene expression of *NaEAH*s and *NaEAS*s in *N. attenuata* after *A. alternata* challenge. Whether or not ethylene is involved in capsidiol biosynthesis is likely dependent on pathosystem. Additionally, our experiments do no indicate a role for jasmonate signaling in the regulation of capsidiol biosynthesis, as transcripts of *NaEAH*s and *NaEAS*s was equivalent in WT and JA deficient irAOC plants at 1 and 3 dpi (Fig. 5).

Despite the great role of capsidiol on resistance, the transcriptional regulation of its biosynthesis is still not clear. Due to the high level of gene expression seen in the capsidiol biosynthetic pathway, we hypothesized that the transcription factors regulating these genes must also be highly expressed during the initial infection period of the fungus. We performed a screen of the 6 most up-regulated transcription factors. Indeed, the NaERF2-like was identified to be a positive regulator of plant resistance and capsidiol production. When compared with EV plants, those with a silenced *NaERF2-like* gene accumulated fewer transcripts of *NaEAS*s and *NaEAH*s, as well as lower levels of capsidiol (Fig. 7). Consistently, *NaERF2-like*-silenced plants were susceptible to the fungus (Fig. 7).

Both *NaEAS* and *NaEAH* are encoded by members of a multi-gene family. Since *NaEAS12* expression is greatly reduced in *NaERF2-like*-silenced plants (Fig. 8), we selected this gene to test whether its promoter could be directly activated by NaERF2-like. Several lines of evidence support that NaERF2-like directly regulates the capsidiol biosynthetic gene *NaEAS12*, including 1) the binding of NaERF2-like protein to the promoter region of *NaEAS12*, which was supported by yeast-one-hybrid, EMSA and Chip-qPCR experiments (Fig. 9); 2) the *NaEAS12* promoter was activated in response to transient *NaERF2-like* over-expression (Fig. 8); this result is further confirmed by stable transgenic lines of Ov-*NaERF2-like*, which exhibit increased *NaEAS12* transcripts and higher levels of capsidiol accumulation (Fig. 10). Currently, it is not known how other *NaEAS*s and *NaEAH*s are regulated by NaERF2-like. Further experiments are needed to test whether or not they are regulated in a way similar to *NaEAS12*.

## Accession Numbers

Sequence data from this article can be found in the GeneBank data library under accession numbers: XM_019375732.1 (HMG-CoA synthase), XM_019375278.1 (MVAPP decarboxylase), XM_019403732.1 (FPP synthase), XM_019409657.1 (Squalene synthase), XM_019408556.1 (EAS12), XM_019399671.1 (NaERF2-like).

## Supplementary data

**Fig. S1. *Alternaria alternata*-elicited *NaEAS12* transcripts in EV and plants individually silenced with the top 5 up-regulated transcription factors.**

Mean (± SE) relative *A. alternata*-induced *NaEAS12* transcripts as measured by real-time PCR in 5 replicated young leaves of EV, VIGS NaERF ABR1-like, VIGS NaZAT12-like, VIGS NaWRKY40, VIGS NaWRKY43 and VIGS NaWRKY61 plants at 3 dpi. Two independent VIGS experiments presented similar results. Asterisks indicate levels of significant differences between EV and VIGS plants (Student’s *t*-test: *, *p*<0.05; **, *p*<0.01; ***, *p*<0.005)

**Fig. S2. Overexpression of NaERF2-like does not affect plant resistance.**

NaERF2-like proteins were detected in 0 leaves of Ov-NaERF2-like line 1 and 2 at 1 dpi by HA antibody via western blot (A). Mean (± SE) diameter of necrotic lesions (B) was recorded in 8-replicated 0 leaves of WT, Ov-NaERF2-like line 1 and 2 plants inoculated with *A. alternata* for 5 d.

**Fig. S3. Silencing *NaEIN2* has a great impact on *A. alternata*-induced transcripts of *NaF6’H1* but does not affect transcripts of *NaEAH*s and *NaEAS*s.**

Mean (± SE) relative *A. alternata*-induced *NaEIN2, NaF6’H1, NaEAS*s, *NaEAH*s transcripts as measured by real-time PCR in 5 replicated young leaves of EV and VIGS NaEIN2 plants at 3 dpi. Asterisks indicate levels of significant differences between EV and VIGS plants with the same treaments (Student’s *t*-test: *, *p*<0.05; **, *p*<0.01; ***, *p*<0.005)

**Table S1.**

Transcriptome analysis revealed regulation of genes involved in sesquiterpene biosynthesis in 3 biological replicate *N. attenuata* leaves after *A. alternata* inoculation at 1 d.

**Table S2.**

Transcriptome analysis revealed top 6 highly elicited transcriptional factor genes in 3 biological replicate *A. alternata*-inoculated *N. attenuata* leaves at 1 d.

**Table S3.**

Primers used in this study.

## Acknowledgements

We thank Prof. Joe Chappell (University of Kentucky, USA) for the capsidiol standard, the Service Center for Experimental Biotechnology of Kunming Institute of Botany, the Chinese Academy of Sciences (CAS), for plant growth support. This project was supported by the National Science Foundation of China (NSFC Grant No. 31670262) and the Key Project of Applied Basic Research Program of Yunnan (2014FA040) to Prof. Jinsong Wu, NSFC grant (No. 31700231), the Applied Basic Research Program of Yunnan (2017FB048) and CAS “Light of West China” Program to Dr. Lan Ma.

## References

Ahuja I, Kissen R, Bones AM. 2012. Phytoalexins in defense against pathogens. Trends in Plant Science, 17, 73–90.

Bailey JA, Burden RS, Vincent GG. 1975. Capsidiol: Antifungal compound produced in Nicotiana tabacum and Nicotiana clevelandii following infection with tobacco necrosis virus. Phytochemistry, 14, 597–597.

Bergelson J, Purrington CB, Palm CJ, Lopez-Gutierrez JC. 1996. Costs of resistance: a test using transgenic Arabidopsis thaliana. Proc Biol Sci, 263, 1659–1663.

Berrocal-Lobo M, Molina A, Solano R. 2002. Constitutive expression of ETHYLENE-RESPONSE-FACTOR1 in Arabidopsis confers resistance to several necrotrophic fungi. The Plant Journal, 29, 23–32.

El Oirdi M, Trapani A, Bouarab K. 2010. The nature of tobacco resistance against Botrytis cinerea depends on the infection structures of the pathogen. Environmental Microbiology, 12, 239–253.

Facchini PJ, Chappell J. 1992. Gene family for an elicitor-induced sesquiterpene cyclase in tobacco. Proceedings of the National Academy of Sciences of the United States of America, 89, 11088–11092.

Glazebrook J. 2005. Contrasting mechanisms of defense against biotrophic and necrotrophic pathogens. Annual Review Phytopathology, 43, 205–227.

Grosskinsky DK, Naseem M, Abdelmohsen UR, Plickert N, Engelke T, Griebel T, Zeier J, Novak O, Strnad M, Pfeifhofer H, van der Graaff E, Simon U, Roitsch T. 2011. Cytokinins mediate resistance against Pseudomonas syringae in tobacco through increased antimicrobial phytoalexin synthesis independent of salicylic acid signaling. Plant Physiology, 157, 815–830.

Guedes MEM, Kuc J, Hammerschmidt R, Bostock R. 1982. Accumulation of 6 sesquiterpenoid phytoalexins in tobacco leaves infiltrated with Pseudomonas lachrymans. Phytochemistry, 21, 2987–2988.

Hao D, Ohme-Takagi M, Sarai A. 1998. Unique mode of GCC box recognition by the DNA-binding domain of ethylene-responsive element-binding factor (ERF domain) in plant. Journal of Biological Chemistry, 273, 26857–26861.

Hao D, Yamasaki K, Sarai A, Ohme-Takagi M. 2002. Determinants in the sequence specific binding of two plant transcription factors, CBF1 and NtERF2, to the DRE and GCC motifs. Biochemistry, 41, 4202–4208.

Huang PY, Catinot J, Zimmerli L. 2016. Ethylene response factors in Arabidopsis immunity. Journal of Experimental Botany, 67, 1231–1241.

Kallenbach M, Bonaventure G, Gilardoni PA, Wissgott A, Baldwin IT. 2012. Empoasca leafhoppers attack wild tobacco plants in a jasmonate-dependent manner and identify jasmonate mutants in natural populations. Proceedings of the National Academy of Sciences, 109, E1548–57.

Kliebenstein DJ, Rowe HC, Denby KJ. 2005. Secondary metabolites influence Arabidopsis/Botrytis interactions: variation in host production and pathogen sensitivity. The Plant Journal, 44, 12.

Krügel T, Lim M, Gase K, Halitschke R, Baldwin IT. 2002. Agrobacterium-mediated transformation of Nicotiana attenuata, a model ecological expression system. Chemoecology, 12, 177–183

LaMondia JA. 2001. Outbreak of brown spot of tobacco caused by Alternaria alternata in Connecticut and Massachusetts. Plant Disease, 85, 230–230.

Li J, Wu J. 2016. Scopolin, a glycoside form of the phytoalexin scopoletin, is likely involved in the resistance of Nicotiana attenuata against Alternaria alternata. Journal of Plant Pathology, 98, 641–644.

Mengiste T. 2012. Plant immunity to necrotrophs. Annual Review of Phytopathology, 50, 267–294.

Mialoundama AS, Heintz D, Debayle D, Rahier A, Camara B, Bouvier F. 2009. Abscisic acid negatively regulates elicitor-induced synthesis of capsidiol in wild tobacco. Plant Physiology, 150, 1556–1566.

Nafisi M, Goregaoker S, Botanga CJ, Glawischnig E, Olsen CE, Halkier BA, Glazebrook J. 2007. Arabidopsis cytochrome P450 monooxygenase 71A13 catalyzes the conversion of indole-3-acetaldoxime in camalexin synthesis. Plant Cell, 19, 2039–2052.

Ohtsu M, Shibata Y, Ojika M, Tamura K, Hara-Nishimura I, Mori H, Kawakita K, Takemoto D. 2014. Nucleoporin 75 is involved in the ethylene-mediated production of phytoalexin for the resistance of Nicotiana benthamiana to Phytophthora infestans. Molecular Plant-Microbe Interactions, 27, 1318–1330.

Pre M, Atallah M, Champion A, De Vos M, Pieterse CMJ, Memelink J. 2008. The AP2/ERF domain transcription factor ORA59 integrates jasmonic acid and ethylene signals in plant defense. Plant Physiology, 147, 1347–1357.

Ralston L, Kwon ST, Schoenbeck M, Ralston J, Schenk DJ, Coates RM, Chappell J. 2001. Cloning, heterologous expression, and functional characterization of 5-epi-aristolochene-1,3-dihydroxylase from tobacco (Nicotiana tabacum). Archives of Biochemistry and Biophysics, 393, 222–235.

Sanchez-Vallet A, Ramos B, Bednarek P, Lopez G, Pislewska-Bednarek M, Schulze-Lefert P, Molina A. 2010. Tryptophan-derived secondary metabolites in Arabidopsis thaliana confer non-host resistance to necrotrophic Plectosphaerella cucumerina fungi. The Plant Journal, 63, 115–127.

Schuck S, Weinhold A, Luu VT, Baldwin IT. 2014. Isolating fungal pathogens from a dynamic disease outbreak in a native plant population to establish plant-pathogen bioassays for the ecological model plant Nicotiana attenuata. PLoS One, 9, e102915.

Shibata Y, Kawakita K, Takemoto D. 2010. Age-related resistance of Nicotiana benthamiana against hemibiotrophic pathogen Phytophthora infestans requires both ethylene- and salicylic acid-mediated signaling pathways. Molecular Plant-Microbe Interactions, 23, 1130–1142.

Shibata Y, Ojika M, Sugiyama A, Yazaki K, Jones DA, Kawakita K, Takemoto D. 2016. The full-size ABCG transporters Nb-ABCG1 and Nb-ABCG2 function in pre- and post-invasion defense against Phytophthora infestans in Nicotiana benthamiana. Plant Cell, 28, 1163–1181.

Solano R, Stepanova A, Chao Q, Ecker JR. 1998. Nuclear events in ethylene signaling: a transcriptional cascade mediated by ETHYLENE-INSENSITIVE3 and ETHYLENE-RESPONSE-FACTOR1. Genes & Development, 12, 3703–3714.

Stoessl A, Unwin CH, Ward EWB. 1972. Postinfectional inhibitors from plants I. Capsidiol, an antifungal compound from capsicum frutescens. Phytopathology, 74, 141–152.

Sun H, Hu X, Ma J, Hettenhausen C, Wang L, Sun G, Wu J, Wu J. 2014a. Requirement of ABA signalling-mediated stomatal closure for resistance of wild tobacco to Alternaria alternata. Plant Pathology, 63, 1070–1077.

Sun H, Wang L, Zhang B, Ma J, Hettenhausen C, Cao G, Sun G, Wu J, Wu J. 2014b. Scopoletin is a phytoalexin against Alternaria alternata in wild tobacco dependent on jasmonate signalling. Journal of Experimental Botany, 65, 4305–4315.

Sun H, Song N, Ma L, Li J, Ma L, Wu J, Wu J. 2017. Ethylene signalling is essential for the resistance of Nicotiana attenuata against Alternaria alternata and phytoalexin scopoletin biosynthesis. Plant Pathology, 66, 277–284.

von Dahl CC, Winz RA, Halitschke R, Kuhnemann F, Gase K, Baldwin IT. 2007. Tuning the herbivore-induced ethylene burst: the role of transcript accumulation and ethylene perception in Nicotiana attenuata. The Plant Journal, 51, 293–307.

Ward EWB, Unwin CH, Stoessl A. 1974. Postinfectional inhibitors from plants 13. Fungitoxicity of phytoalexin, capsidiol, and related sesquiterpenes. Canadian Journal of Botany-Revue Canadienne De Botanique, 52, 2481–2488.

Welsch R, Maass D, Voegel T, Dellapenna D, Beyer P. 2007. Transcription factor RAP2.2 and its interacting partner SINAT2: stable elements in the carotenogenesis of Arabidopsis leaves. Plant Physiology, 145, 1073–1085.

Wu J, Wang L, Wunsche H, Baldwin IT. 2013. Narboh D, a respiratory burst oxidase homolog in Nicotiana attenuata, is required for late defense responses after herbivore attack. Journal of Integrative Plant Biology, 55, 187–198.

Xu Z, Song N, Ma L, Fang D, Wu J. 2018. NaPDR1 and NaPDR1-like are essential for Nicotiana attenuata resistance to the fungal pathogen Alternaria alternata. Plant Diversity, 40, 68–73.

Yoo SD, Cho YH, Sheen J. 2007. Arabidopsis mesophyll protoplasts: a versatile cell system for transient gene expression analysis. Nature Protocols, 2, 1565–1572.

Zhao Y, Wei T, Yin KQ, Chen Z, Gu H, Qu LJ, Qin G. 2012. Arabidopsis RAP2.2 plays an important role in plant resistance to Botrytis cinerea and ethylene responses. New Phytologist, 195, 450–460.

